# The multilevel society of proboscis monkeys with a possible patrilineal basis

**DOI:** 10.1101/2023.01.24.525467

**Authors:** Ikki Matsuda, Tadahiro Murai, Cyril C. Grueter, Augustine Tuuga, Benoit Goossens, Henry Bernard, Nurhartini Kamalia Yahya, Pablo Orozco-terWengel, Milena Salgado-Lynn

**Author notes:** Co.First Author (Ikki Matsuda, Tadahiro Murai).

## Abstract

Multilevel societies (MLS), which are characterized by two or more levels of social organization, are among the most complex primate social systems. MLS have only been recorded in a limited number of primates, including humans. The aim of this study was to investigate whether proboscis monkeys (*Nasalis larvatus*) form MLS in Sabah, Malaysia, and to genetically characterize their dispersal patterns. Association data were obtained through direct observation (35 months) and kinship data through genetic analysis, based on feces collected from ∼200 individuals. The results strongly suggest that proboscis monkeys exhibit a form of MLS, with several core reproductive units and a bachelor group woven together into a higher-level band. Genetic analysis revealed that the females migrated randomly over short and long distances; however, the males tended to migrate relatively shorter distances than females. Furthermore, male-male dyads showed a slightly higher average relatedness than female-female dyads. Combined with the results of direct observations, we conclude that proboscis monkeys form a MLS with at least two layers and a patrilineal basis. Since patrilineal MLS have been identified as an important step in the evolution of human societies, their convergent appearance in proboscis monkeys may help us understand of the drivers of human social evolution.

**Significance Statement:** The aim of this study was to determine the social organization of proboscis monkeys by direct observation and genetic analysis. The results revealed that their social system exhibited a form of multilevel society with a possible patrilineal basis. Since humans exhibit a similar constellation of social features, proboscis monkeys may offer insightful clues about human social evolution.

## Introduction

Primate multilevel societies (MLS) are a form of social organization where the joint activities of a set of core social units (foraging, resting, and traveling) scale-up to generate higher societal levels, such as bands (Grueter et al. 2020). Units do not have territorial boundaries, but ranges, which completely overlap with those of other units. Interactions with other units are characterized by a lack of persistent hostility, with units generally tolerating each other’s presence. Although the units are part of a larger collective (e.g., a band), they can temporarily bud off from the band (e.g., for foraging). This form of fission–fusion where the integrity of the core unit is maintained, is distinct from the more atomistic pattern seen in single-tiered societies, such as those of chimpanzees (*Pan troglodytes*) and spider monkeys (*Ateles* spp.), where a group (community) divides into smaller sub-groups (parties) of variable size or composition and each community has its own territory or home range (Grueter et al. 2012a; Rodseth et al. 1991).

A prime example of a species where MLS are almost universally present is humans (Dyble et al. 2016; Grueter et al. 2012a; Rodseth et al. 1991). Human MLS are based on family groups that are integrated into a network of increasingly larger social tiers, such as bands and tribes (Layton and O’Hara 2010). The MLS of nonhuman primates are not as complex as human MLS. However, several tiers of social stratification can be found in a limited number of species (Grueter et al. 2012b). MLS have evolved in African papionin species, such as geladas (*Theropithecus gelada*) and hamadryas baboons (*Papio hamadryas*) (Dunbar and Dunbar 1975; Kummer 1968; Swedell 2011; Swedell and Plummer 2012), and in Asian colobine species, in particular, the odd-nosed group, including snub-nosed (*Rhinopithecus* spp.) and proboscis monkeys (*Nasalis larvatus*) (Grueter et al. 2022). The minimum reproductive unit in the MLS of most nonhuman primates is the one-male–multifemale group, but MLS that are not based on such groups, i.e., multimale–multifemale groups, have recently been reported in Guinea baboons (*Papio papio*) (Patzelt et al. 2014, but also see Goffe. et al. 2016) and Rwenzori colobus (*Colobus angolensis ruwenzorii*) (Miller et al. 2020; Stead and Teichroeb 2019).

This nested arrangement of grouping tiers allows individuals to be concurrently associated with multiple grouping levels and to harness the adaptive benefits that each level entails while minimizing costs such as feeding competition (Grueter et al. 2017). For example, the band level allows core units to ‘recruit’ extra-unit males for collective defense when needed (Xiang et al. 2013) and the core unit offers a safe haven for individuals to exchange social services, such as grooming or allocare (e.g. Dunbar 1979; Yu et al. 2013). When fission–fusion of core units is superimposed on a multilevel organization, the system becomes even more flexible, as core units also have the option to separate from the higher-level group when the availability and distribution of food resources demands it (Schreier and Swedell 2012).

By studying the characteristics of MLS in distantly related species we can understand what environmental conditions and/or potential selection pressures are associated with such traits in different lineages. MLS are generally absent in the primates most closely related to humans (i.e., the great apes), although evidence is inconclusive in some cases (Grueter and Wilson 2021; but see Morrison et al. 2019). Consequently, studies on primates that are phylogenetically distantly related but exhibit multilevel structures are important for elucidating the evolutionary paths leading to MLS in humans (Grueter et al. 2012a).

Proboscis monkeys are endemic to the island of Borneo. This species is a large, sexually dimorphic arboreal colobine that typically inhabits mangrove forests, peat swamps, and riverine forests (Grueter et al. 2022). The basic components of their society, i.e., the minimum reproductive units (core units), are one-male–multifemale units (OMUs) which assemble with each other in riverside trees (Bennett and Sebastian 1988; Boonratana 2002; Yeager 1991), with varying degrees of aggregation depending on seasonal fluctuations in food abundance or predation pressure (Matsuda et al. 2010a). Several studies suggest that there is a multilayered social organization to this association of OMUs along the river (Bennett and Sebastian 1988; Boonratana 2002; Yeager 1991), although one study suggests that ecological factors, such as food availability and predation pressure, would be sufficient to explain the cohesion of OMUs without assuming a higher order structure of this type (Matsuda et al. 2010a). In addition to OMUs, all-male or bachelor groups (AMGs) are also found, consisting mostly of immature males who are often accompanied by adult males (Matsuda et al. 2020a; Murai 2004). Although males of these AMGs potentially start OMUs in the future (Murai 2004), little is known about their social structuring and no study yet has examined the social structure of proboscis monkeys from a genetic basis.

The aim of this study was to determine the social structure of proboscis monkeys both by direct observation of multiple identified groups and genetic analysis, which was not achieved in previous studies (Boonratana 2002; Matsuda et al. 2010a). The six-month study of Yeager was based on the identification of multiple groups (OMUs and AMGs) and examining the interactions between them (Yeager 1991). However, we observed long-term interactions between the groups for 35 months. Given that proboscis monkeys are thought to form MLS, certain OMUs are expected to frequently maintain close proximity on their sleeping trees along the river. To gain a comprehensive picture of their complex society, we analyzed the degree of kinship between individuals within a population of proboscis monkeys through fecal DNA, albeit at a different time of the year from the time of direct observation. In proboscis monkeys, both males and females disperse from their natal OMUs to other OMUs or AMGs before they reach maturity (Matsuda et al. 2012a). However, their social networks, based on grooming interactions, are decidedly female-centered (Matsuda et al. 2012b; Yeager 1990b). We assume that the emergence of societies in which females are central to social interactions would be based on a high degree of kinship between females within OMUs (Tinsley Johnson et al. 2013). Therefore, we hypothesize that although both sexes disperse between groups, females would tend to disperse to more proximal OMUs than males (Guo et al. 2015, but see Fang et al. 2022) and/or to be motivated to maintain post-dispersal contact with maternal female kin (Städele et al. 2016). We also hypothesize that males disperse over longer distances than females, whose dispersal distances are shorter. Lastly, we discuss the similarities and differences between the findings from this study and those from studies of closely related snub-nosed monkeys, with better studied MLS (Grueter 2022).

## Methods

### Study site and animals

We conducted the study in a riverine forest along the Menanggul River, a tributary of the Kinabatangan River, Sabah, Malaysia (118°30′ E, 5°30′ N). The mean minimum and maximum daily temperatures were approximately 24°C and 30°C, respectively, and the mean annual precipitation at the site was 2,474 mm (Matsuda et al. 2019). The river level varied by approximately 1 m daily. The average river level rose >3 m during seasonal flooding. The riverine forest was inhabited by eight diurnal primate species, including our study subject, the proboscis monkey. Proboscis monkeys under observation were well habituated to observers in boats, because this area is one of the main tourist attractions in the region, with many boats and tourists visiting the Menanggul River almost on a daily basis during the last decade.

### Behavioral observation in 1999–2002

Along the Menanggul River, using a GPS unit, location points were collected at intervals of 50 m over a transect stretching from the river mouth of the Menanggul River up to a point 6 km upstream. Proboscis monkeys typically return at night to the riverbank to sleep in our study site (Matsuda et al. 2010b). In this study, the core observation area was set up from the river mouth up to 4 km upstream, where boat-based observations were performed in the early morning (06:00–09:00) and/or late afternoon (15:00–18:00). However, if we could not find groups of proboscis monkeys in the area for up to 4 km, we extended the survey to the upstream area of up to 6 km.

We conducted the behavioral observations from February 1999 to October 2001, and in May and June 2002 (560 d). By late June 1999, eight OMUs and one AMG were identified by distinguishing all adult males and a few adult females based on their physical features, such as scars and nose shapes. The number of OMUs observed in the study area was consistently eight (Koda et al. 2018); the mean number of observed days for each group (OMUs and AMGs) was 187.3 (standard deviation = 117.5) with a range from 12 to 366 days (see Supplementary Table 1 in details). The replacement of the dominant males, named Ba, Wa, and Bu, was observed on three occasions during the study period in August 2000, November 2000, and October 2001, respectively (Murai et al. 2007). In each case, the replacing males came from the AMGs. We refer to OMUs before the male replacement occurred as “senior” OMUs (Ts, Be, Po, Bo, Yo, De, Pu, and Ki) and OMUs after the male replacement as “junior” OMUs (BoWa, PoBu, and KiBa) for the sake of convenience. Throughout the study period, the mean number of individuals comprising OMUs was 18, ranging from 8 to 34, and AMGs consisted of approximately 30 males (Supplementary Table 2). The names of the OMUs and AMGs and the locations of their sleeping sites were recorded whenever proboscis monkeys were found along the river in the late afternoon.

### Genetic analysis

#### DNA sampling in 2015–2016

From July 2015 to April 2016, we collected fecal samples from proboscis monkeys in the study area for genetic analyses. A boat survey was conducted in the late afternoon to detect the group locations and record the GPS coordinates of their sleeping sites: early morning the next day, when the monkeys were still asleep, the sites were revisited while we waited in the boat until the monkeys moved inside the forest. As proboscis monkeys often defecated shortly before moving into the forest, fresh feces were collected by carefully exploring the ground near their sleeping trees after they had left them. However, several groups often stayed in the trees along the river in close proximity and it was difficult to reliably distinguish the group to which the respective feces on the forest floor corresponded. We rubbed the surface of the fecal pellets with cotton swabs, and dipped the swabs in 2 mL tubes containing 1 mL lysis buffer, consisting of 0.5% SDS, 100 mM EDTA, 100 mM Tris-HCl, and 10 mM NaCl (Longmire et al. 1997). The swabs were discarded, and the tubes were stored at room temperature for ∼400 d after collection until they were delivered to the laboratory, where they were kept at −80°C until DNA extraction. The DNA of 307 samples was extracted with QIAamp DNA Stool Mini Kit (Qiagen) using TE (1 M Tris-Cl, 0.05 M EDTA; pH 8.0) as elution buffer.

### Microsatellite screening and sex identification

Twenty-four markers (23 polymorphic microsatellite loci and 1 fragment of the DEAD-box gene) were amplified for genetic analyses. The DEAD-box gene was used to identify sex (Inoue *et al*. 2016). The microsatellite loci were selected based on the method of Inoue *et al*. (2016) and Salgado-Lynn *et al*. (2010a), with four PCR multiplex reactions optimized for this study (Supplementary Table 2).

PCR was performed using the Applied Biosystems Veriti 96 Well Thermal Cycler PCR machine. All PCR amplifications were performed in 15 μL reactions, containing 1 × Master Mix (Qiagen Multiplex Kit), 0.4 μg/μL of BSA, 0.2 μM of each primer used for the multiplex combinations, and 2 μL of 10–100 ng of template DNA. After an initial incubation at 95°C for 15 min, PCR amplification was performed for 40 cycles, consisting of denaturation at 94°C for 45 s, annealing at 60°C or 62°C for 90 s, extension at 72°C for 90 s, and a final extension at 72°C for 10 min. Then, 2% agarose gels were used to visualize band quality and verify target band size. The PCR products were sent to First BASE Laboratories (Kuala Lumpur, Malaysia) for fragment analysis. The results of fragment analysis were scored using the software GeneMarker version 2.6.3 and corrected by eye.

We used 20 samples for a pilot study to determine the number of positive PCR repetitions needed to obtain a reliable genotype. A consensus threshold (100 simulations; the range of repetitions was from two to several) was produced in GEMINI v.1.4.1 (Valière *et al*., 2002) in the “Consensus Threshold Test” module. The consensus threshold was used in a “PCR Repetition Test,” also in GEMINI (1,000 simulations; the range of repetition was two to 10), which showed that three positive repetitions for each multiplex were sufficient to achieve a reliable genotype, with a maximum of five positive repetitions to clarify ambiguities. However, if there were no PCR products after three PCR rounds in two or more multiplexes, the sample was not included in the analysis.

Of the 307 samples from which DNA extraction was performed, 267 fecal samples were genotyped, and only 197 were included in relatedness analysis (<10% missing data). Noted that feces from the same individuals were excluded to prevent duplicate analyses after the genetic profiling; a sample of the same individual with the least missing data was retained for analysis. GenAlEx 6.51b2 (Peakall and Smouse, 2006, 2012) was used to calculate heterozygosity, frequency-based statistics, and polymorphism (see Supplementary Table 3). A Hardy–Weinberg Equilibrium (HWE) exact test was performed separately for males (N = 64) and females (N = 133) in Genepop 4.7.5 (Raymond and Rousset 1995; Rousset 2008) using default Markov chain parameter values. Neither group was in HWE (*p* < 0.001).

### Data analysis

#### Degree of association between groups

The distance between the edges of the sleeping site location of each OMU or AMG along the river was used as a criterion to determine if they were alone or in association. In association was defined as 100 m or less between the edges of the sleeping site of the relevant OMU or AMG. The rationale for this approach follows previous studies where OMUs of few individuals or OMUs with identified or unidentified individuals were considered members of an OMU if they were observed to be within a distance of 100 m of an OMU (Kawabe and Mano 1972; Kern 1964; Macdonald 1982; Salter et al. 1985). Indeed, long-term focal tracking of a specific OMU also indicates that group members are rarely more than 50 m from each other (Matsuda et al. 2009), hence it would be appropriate to define the close proximity of two or more groups as within 100 m of each other, which was also recommended as being operationally suitable by Yeager (1992). As a result, OMUs were in close proximity within 100 m in 23.7% of the total number of observation days in this study (Supplementary figure 1). However, the frequency with which the relevant groups had sleeping sites within 100 m from each other is not an appropriate index of the degree of their association as the frequency of appearance of each group along the river varied. Thus, to assess intergroup associations, we calculated the percentage of days when the relevant groups had sleeping sites within 100 m from each other, based on the number of days when the sleeping sites of relevant groups were detected along the river. As all groups were identifiable and boat surveys along the riverside were carried out one way, no duplicate groups were observed during the survey time. In addition, although groups almost always return to the riverside trees in the late afternoon to sleep (Matsuda et al. 2010b), each group appeared along the riverside at a different time of day. Therefore, it was not always spotted them all during the survey time.

We generated weighted social networks between the groups based on the degree of association calculated using the R package ‘igraph’ (Csardi and Nepusz 2006). In this study, two types of social network diagrams were drawn including all study groups to visualise how their social network changed, before and after the replacement of the dominant male in the OMGs (see detailed observation periods of each group in Supplementary Table 1), i.e., the network including the data from 8 senior OMUs and 1 AMG (before replacement) and the one including the data from 5 senior OMUs, 3 junior OMUs and 1 AMG (after replacement). Additionally, we used cluster analysis (Quinn and Keough 2002) to assess similarity relationships for the associations among study groups based on the same dataset using the social networks. The degree of association among groups was clustered using Euclidean distance and Ward’s linkage method (Quinn and Keough 2002). The cluster analysis did not include the OMU “PoBu” due to limited observation cases, i.e. < 20 caese (see Supplementary Table 1 for details). The quality of a clustering was validated by an analysis of silhouette values (Rousseeuw 1987), which vary from −1 to 1 and represent the tightness of the data points within a cluster and the separation between different clusters in a given model. The silhouette value *S*(*x*_*i*_) for a single data point *x*_*i*_ is computed as

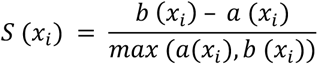

where *a* (*x*_*i*_) is the average Euclidean distance of *x*_*i*_ to all points in the same cluster, and *b* (*x*_*i*_) is the minimum average Euclidean distance of *x*_*i*_ to all other clusters in which *x*_*i*_ is not a member. Thus, low values of *a* (*x*_*i*_) indicate that *x*_*i*_ is representative of the cluster, whereas high values of *b* (*x*_*i*_) indicate that *x*_*i*_ is much different from the other clusters. Higher values denote higher clustering quality, whereas negative values denote that the data point should otherwise belong to a different cluster. By computing,

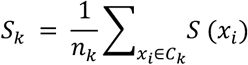

as the average of *S*(*x*_*i*_) over all *n*_*k*_ points in cluster *C*_*k*_, one obtains an indicator of how well separated the cluster is from all others. An indicator for the complete clustering is obtained by averaging *S*_*k*_ over all *K* deduces both *a*(*x*_*i*_) and *b*(*x*_*i*_) simultaneously, and thus, a larger number of clusters does not necessarily improve the silhouette score. This property makes the silhouette a better indicator for cluster quality than, for example, the log-likelihood of data, because the latter merely increase with a larger *K*.

#### Relatedness and Fst

We used the Moment Estimate of Relatedness method of Wang (2002) as implemented in Mer2 to estimate relatedness between all pairs of individuals and to calculate mean total relatedness (R) and its standard deviation using 10,000 bootstrap replicates, which simulate allele frequencies. Using a Kolmogorov–Smirnov test (KS-test), we tested for differences in relatedness between male-male and female-female dyads in two data sets. The first data set included all dyads, and the values for dyads with negative estimated relatedness estimates were transformed to zero as negative relatedness values indicate no shared alleles due to common descent.

The effect of spatial autocorrelation was measured to determine the effect of geographical distance between individuals and the relatedness they showed for females and males separately. For this, we used the geographic distance between individuals measured with respect to the mouth of the river and R values between them with negative values converted to zero (i.e., no kinship). Based on these data, we tested whether there was a correlation between geographical distance and genetic distance (isolation by distance) using the Kendall rank correlation coefficient as the data was not normally distributed (one sample KS-test *p*-value = 2.2e-16).

Given that our analysis counted each sample as a single point, the statistical power to detect a relationship between degree of relatedness and geographical distance was potentially weak. To increase the statistical power of the data, we subdivided the entire set of samples into zones (i.e., with multiple pairwise comparisons per zone to more accurately estimate means and variances) and tested for a relationship between the mean relatedness between zones and the corresponding geographic distance. In other words, the rationale for carrying out this analysis using bins is that by grouping samples in bins we increased the statistical power to detect differences between bins. Based on the location points along the river where the fecal samples were collected, we divided the locations into distance bins as measured from the river mouth: i) 0–500, ii) 501–1000, iii) 1501–2000, iv) 2001–2500, v) 2501–3000, vi) 3001–3500, and vii) 3501–4000 m. Notably, we excluded zone 1001–1500 m, because we collected only one fecal sample in that section. For each distance bin we calculated the average relatedness of the dyads in the bin and used that for the correlation against geographical bins and the upper boundary of the distance bin. Based on these data, we tested whether there was a correlation between geographical zone and genetic distance using the Pearson’s *r* was used as the data was normally distributed (Shapiro Wilk test *p*-value>0.05). In addition, within those sampling zones, we estimated pairwise divergence *F*_*ST*_ of Wright (1951) with the method of Weir and Cockerham (1984) between all pairs of zones and we used Mantel tests to determine the relationship of the matrix of geographical distance between pairs of zones, e.g., two adjacent zones were separated by 500 m, whereas the most distant zones were separated by 4000 m. The *p*-value threshold for the Mantel test was corrected with the FDR method of Benjamini and Yekutieli (2001). The *F*_*ST*_ range is from zero (no differences between allele frequencies in the two groups compared) to one (completely different allele frequencies in the two groups compared). We calculated them with the package hierfstat (Goudet 2005) in R (R-Core-Development-Team 2023).

#### Limitations of analysing data collected at different time periods

In this study, although behavioural data were collected by direct observation based on individually identified groups, genetic samples were collected from the same population more than 10 years after those observations were carried out, and the data collected at these different times were combined to discuss the social structure of the proboscis monkeys. When the genetic samples were collected, the previous identification sheets could not be applied, so the samples had to be collected without identifying individuals. Noted, however, even if a group could be identified, as noted above, several groups often stayed in close proximity to trees along the river, making it difficult to reliably distinguish each faeces on the forest floor corresponding to which group. Anyhow, although the data were collected from the same population, it should be noted in interpreting the results of this study that the behavioural observation data and the genetic data were not linked in a real-time context. However, we believe that even if generational changes in individuals would occur over a period of 10 years, the overall basic social structure would not have changed significantly and the genetic structure within the population would not have changed significantly either. This is because the population dynamics of the proboscis monkeys at the study site remained stable within the 10-year period (Matsuda et al. 2020a).

## Results

### Direct observation of social structure in 1999–2002

The total number of observations per group is provided in Supplementary Table 1. Groups differed in the number of observations, ranging from 12 (PoBu) to 366 (Po). The weighted graph, based on the degree of association of each group pair, is shown in Figure 1a. Based on the evaluation using silhouette values, hierarchical cluster analysis indicated two major clusters, i.e., band A, including the OMUs, Ts, Be, Po, and Bo (*S*_*k*_ = 0.236); and band B, including the OMUs, Yo, De, Pu, and Ki, and the AMG (*S*_*k*_ = 0.273; Figure 1b). A junior OMU (BoWa) kept belonging to band A (*S*_*k*_ = 0.186) even after the group was replaced by young males (Wa) from the AMG (Figure 1c). Replaced PoBu was not included to the clustering analysis due to the limited number of observed cases, i.e. < 20. However, another junior OMU (KiBa) moved to band A from band B (*S*_*k*_ = 0.308) after the group of Ki was replaced by Ba. The AMG flexibly changed the band while replacements occurred (Figure 1d).

**Figure 1.**
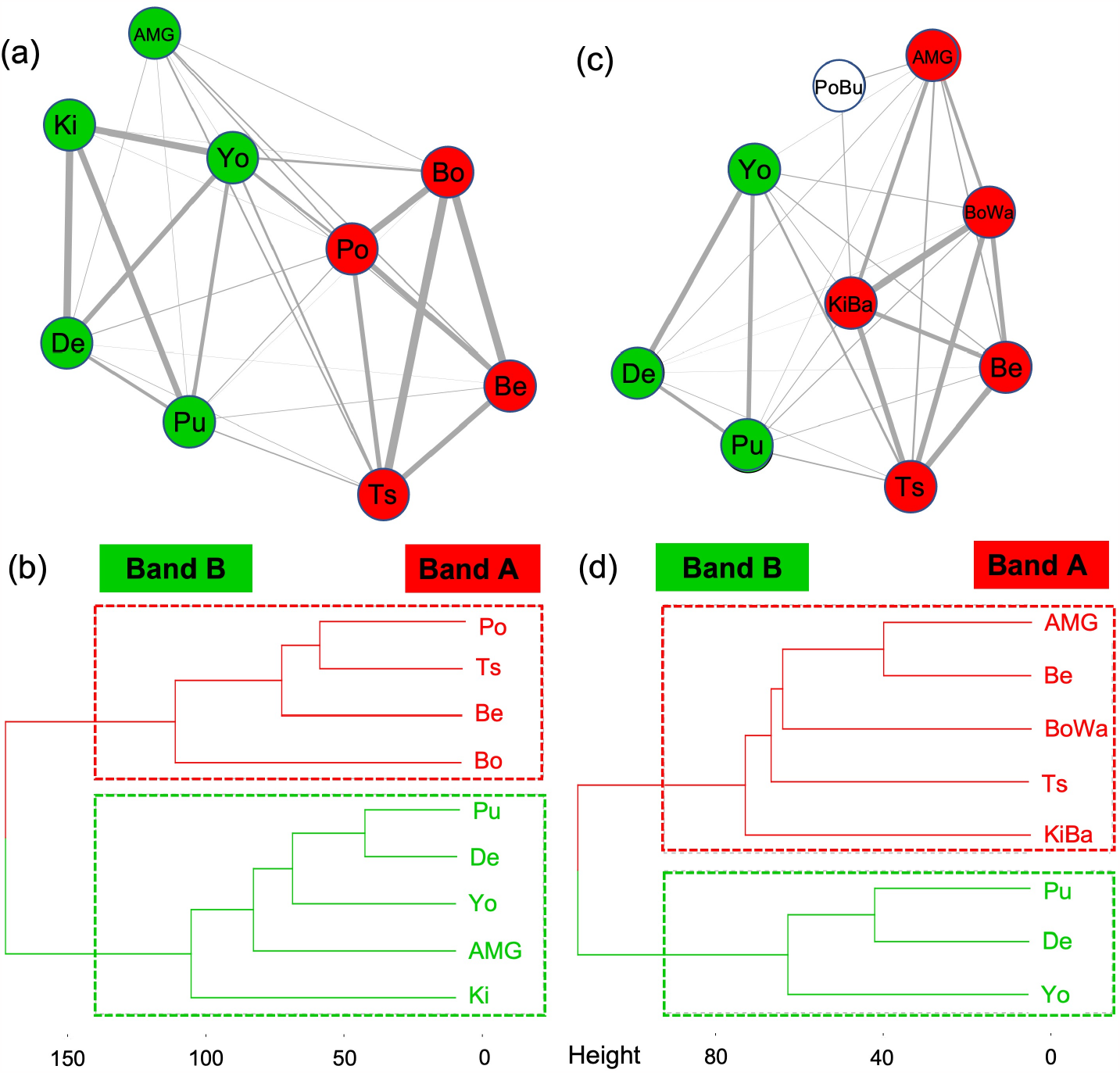
Network graph of the OMUs and AMG identified with cluster analyses, illustrated based on the association index. The thickness of the edge reflects the degree of association index and the height between two clusters in a dendrogram indicates the dissimilarity/distance between them, from February 1999 to August 2000, when the first replacement of males in OMUs was observed (a, b), and thereafter, to June 2002, including two further replacements of males in OMUs (c, d). The cluster analysis did not include PoBu due to limited observation cases, i.e. < 20, and the two statistically estimated communities (bands) are colored.

The two bands in the study site were likely to segregate to areas in which they slept along the Menanggul River, although these widely overlapped. The upstream area of the Menanggul River was predominantly occupied by band A, as shown by a mean distance of sleeping sites from the river mouth of 923 m, ranging from 0 m to 2000 m (Figure 2). By contrast, band B mostly occupied the downstream area, with a mean sleeping site distance from the river mouth of 2,042 m, ranging between 400 and 5,800 m. The home range of the AMG spread over the Menanggul River, with a mean sleeping site distance from the mouth of the Menanggul of 2,837 m and a wider range from 0 m to 6,000 m.

**Figure 2.**
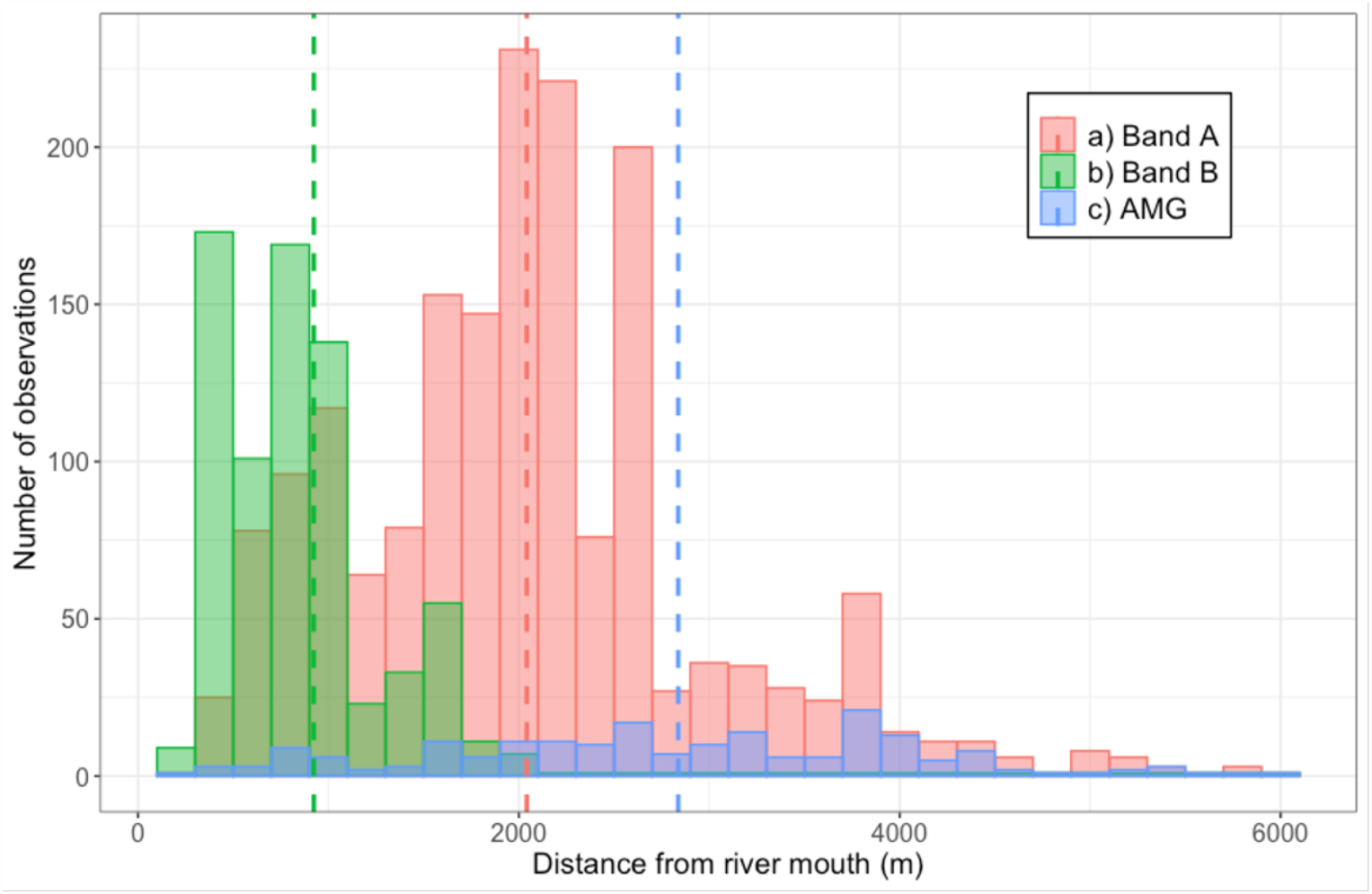
Distribution of sleeping sites of each band and all-male group (AMG) along the Menanggul River. The dotted lines in the graph represent the mean location of the sleeping location based on the distance from the river mouth of each band and AMG. Different bands and the AMG are colored.

### Genetic analysis of social structure in 2015–2016

#### Overall relatedness pattern

The total number of relatedness pairwise comparisons measured among the 197 samples was 35,511, of which 27,777 (78%) were negative numbers, thus indicating that those pairs of samples are not related to each other. Of the remaining dyads with positive relatedness values, there were no identical twins, 187 parent-offspring or full sibling dyads, and ∼1207 relationships that could be considered cousins or half-siblings (Supplementary Figure 2).

#### Female vs. male relatedness

There was a significant difference in relatedness values between male-male and female-female dyads when including all same-sex dyads (KS-test *p*-value = 4.89e-6), with the males showing on average a higher relatedness (0.0481 ± 0.109) than the females (0.0369 ± 0.094).

#### Relatedness based spatial autocorrelation

Based on relatedness values, a significant negative correlation between geographic distance and genetic distance was found for both sexes, with females (N = 9045 dyads) showing a negative Kendall’s τ = −0.066 (*p*-value = 1.153e-15) and males (N = 2016 dyads) showing a slightly less negative Kendall’s τ = −0.053 (*p*-value = 0.00153; Figure 3). Overall, as geographic distance increased, the degree of relatedness decreased, indicating that the more distant the two individuals were, the lower their relatedness.

**Figure 3.**
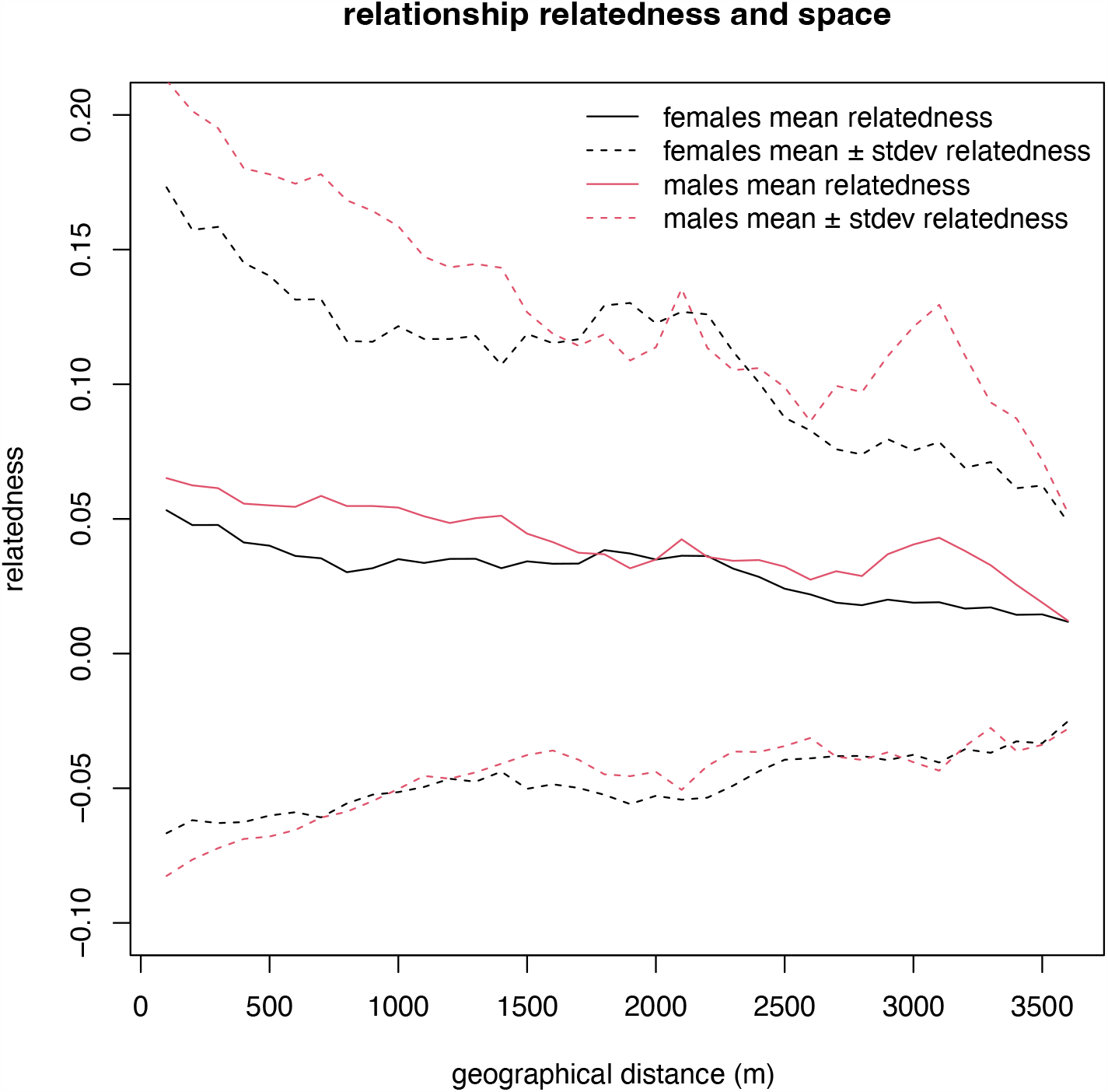
Relationship between geographic distance and genetic distance among the 197 samples measured. The solid lines are the mean relatedness across distance classes for the males (red) and females (black) and the dashed lines are the upper and lower boundaries of the mean plus and minus standard deviation of the relatedness for males and females across distances, respectively.

When investigating the correlation between relatedness values and distance zone, for each sex, a significant reduction was observed in relatedness as geographical distance increased (female Pearson’s *r* = −0.906, *p*-value = 0.0048; male Pearson’s *r =* −0.947, *p*-value = 0.001182), with approximately 82% of the difference in relatedness explained by distance for females and 89% for males.

#### Pairwise differentiation along the river and isolation by distance

The relationship between geographical distance and degree of differentiation (*F*_*ST*_) was examined after dividing the samples into zones based on the location from which their fecal samples were collected. Significant differences were found in the degree of *F*_*ST*_ in the pairwise comparisons of 4 out of 21 zones-by-zone comparisons for males (Table 1). Males tended to exhibit significant differentiation with individuals living in the uppermost zone of the river (3501–4000 m) from those living near the river mouth, which is indicated by an increase in genetic differentiation with an increase in geographical space between them (Mantel test squared correlation coefficient r^2^ = 0.603, *p*-value = 0.0055), and thus, approximately 60% of the difference in allele frequency between males can be explained by the geographical distance between the sampling points. Although females in 5 out of 21 zones-by-zone comparisons showed significant genetic differentiation based on their FST values (Table 1), FST values did not increase with geographic distance (Mantel test squared correlation coefficient *r*^*2*^ = 0.022, *p*-value = 0.67302).

**Table 1.** Pairwise differentiation (*F*_*ST*_) along the river, divided the locations into distance bins that reflected samples separated from each other by i) 0–500, ii) 501–1000, iii) 1501–2000, iv) 2001–2500, v) 2501–3000, vi) 3001–3500, and vii) 3501–4000 m. Notably, the zone 1001–1500 m was excluded as only one fecal sample was collected in that section. Bold numbers denote the pairs that were statistically significant.

|  | 0-500m | 501-1000m | 1501-2000m | 2001-2500m | 2501-3000m | 3001-3500m | 3501-4000m |
| --- | --- | --- | --- | --- | --- | --- | --- |
|  | Female |  |  |  |  |  |  |
| 0-500m |  | 0.00850 | **0.02932** | 0.01433 | **0.03762** | 0.02474 | 0.02557 |
| 501-1000m | 0.01416 |  | 0.00895 | 0.01453 | 0.02103 | 0.01285 | 0.00865 |
| 1501-2000m | −0.02457 | −0.04066 |  | 0.01333 | **0.03910** | **0.02815** | 0.02389 |
| 2001-2500m | 0.00955 | 0.02545 | 0.00496 |  | **0.03190** | 0.02033 | 0.01077 |
| 2501-3000m | 0.02383 | −0.00277 | −0.02280 | **0.02912** |  | 0.03837 | −0.01313 |
| 3001-3500m | 0.04369 | 0.01264 | 0.03425 | 0.02159 | 0.02017 |  | 0.00115 |
| 3501-4000m | **0.05916** | **0.05027** | **0.08786** | 0.04359 | 0.02889 | 0.03311 |  |

## Discussion

The findings strongly suggest that the proboscis monkey exhibits a form of MLS, consisting of several core reproductive units and bachelor groups interwoven with higher-order bands. Our study also highlights sex-specific relatedness differences, a negative correlation between relatedness and geographic distance, and distinct genetic differentiation patterns along the river for males and females. Notably, males showed slightly higher relatedness than females, though a negative correlation between relatedness and geographic distance was consistent for both sexes, indicating decreasing relatedness with increased distance. In addition, pairwise differentiation along the river was significant for males, with 60% of allele frequency differences explained by geographic distance, though females showed genetic differentiation in specific zones without an increase in differentiation with distance. These results suggest that females migrated randomly across both short and long distances, whereas males tended to migrate relatively shorter distances than females, possibly leading to patrilineal characteristics within the population of the studied species. Thus, it is more likely that our hypothesis that proboscis monkeys form a matrilineal, multileveled society is ruled out and that it would be rather patrilineal. The form of MLS inferred from either the results of direct observation or genetic analysis, and that inferred from the integration of these two results, will be discussed in more detail below.

### Inferring MLS based on observational data

The findings of this study revealed that the social system of the proboscis monkey takes the form of a MLS, with OMUs (core reproductive units) and the AMG (bachelor group) woven together into a higher-level band. Several studies have reported that OMUs of proboscis monkeys are not territorial and sleep in close proximity in the trees along the riverside where they spend the night (Bennett and Sebastian 1988; Boonratana 2002). In particular, the study by Yeager (1991), in which multiple groups were identified and their relationships examined over six months, provides the most detailed analysis of the social organization of proboscis monkeys. The results of Yeager are generally in accordance with our results: two bands consisting of multiple OMUs and AMGs were detected, and the two bands were loosely segregated upstream and downstream of the river, with a certain degree of overlap in their distribution ranges.

Although this study confirmed that the proboscis monkey forms MLS, a previous study conducted by us at the same site (Matsuda et al. 2010a) was ambiguous in this regard. That study showed that changes in local density of OMUs along the river (defined as the number of observed sleeping sites of OMUs other than the focal OMU on a 1,000 m long segment along the river using the sleeping site of the focal OMU as the midpoint) can be predicted by spatial heterogeneity along the river in relation to predation pressure and food availability. Namely, locations with a narrower river width are advantageous for predator avoidance, because animals can escape by leaping to the other side of the river when attacked by a predator, and better foraging locations where clumped food patches are abundant are also advantageous. In other words, the local density of OMUs may increase where better sleeping conditions are available, which suggests that associations among units are induced by habitat features and are not necessarily the outcome of social attraction. However, this previous study focused on only one identified focal OMU, whereas other OMUs that stayed around that focal OMU were not identified and treated as local density. Hence, the aspect of whether a particular affinity among specific OMUs existed during the period of increased local density was ignored. Integrating the results of our previous study and this study, it appears that the proboscis monkeys form a MLS with specific OMUs at their core, and that the degree of association among units varies seasonally, depending on food abundance and predation pressure. Indeed, the association between identified OMUs of proboscis monkeys at other study sites have been reported to be seasonal, although no ecological explanations for such seasonality were provided (Yeager 1991). A detailed review has shown that ecological conditions, such as food abundance, are merely permissive and do not seem to drive the nested nature of the MLS of Asian colobines (Grueter and van Schaik 2009). However, it is premature to discard the impact of ecological factors, such as resource abundance and predation pressure, on the evolution of MLS in proboscis monkeys (see also Grueter 2022).

In our longer-term study of 35 months, we observed three occasions of adult male replacement in the OMUs, which offered new insights into the social system of the proboscis monkey. We found that, after the replacement of the dominant male, an OMU would occasionally switch band affiliation. Changes in band affiliation have also been documented for the AMG. Our observational protocol did not allow us to explore the reasons behind switches in band affiliation. These could include avoidance of social disadvantages, e.g., a low unit dominance rank. Understanding the motivations for changes in band affiliation requires further detailed individual-level observations of both OMUs and AMG.

### Inferring MLS based on genetic data

Although several genetic studies have been performed on proboscis monkeys, e.g., population genetics focusing on mitochondrial control region (Munshi-South and Bernard 2011), development of polymorphic microsatellite loci (Inoue et al. 2016; Salgado-Lynn et al. 2010), and their application to zoo studies (Ogata and Seino 2014), this study was the first to examine their social structure genetically by estimating relatedness in a large number of individuals (N=197) in the wild. Nonetheless, the results still need to be interpreted with caution due to the potential error-prone of estimating dyadic relatedness from a limited number of microsatellites. Using 24 markers still carries a chance of such a misclassification of kinship (Csillery et al. 2006; Van Horn et al. 2008). In fact, the standard deviation of the estimated relatedness values was large.

The negative correlation between genetic and geographic distances is comparable to that reported for many other primates (e.g. Hagell et al. 2013; Mbora and McPeek 2014; Oklander et al. 2017). By contrast, the significantly higher degree of relatedness between males than females in the population at the study site was rather unexpected in the proboscis monkey, for which a more female-centered social interaction network has been observed (Matsuda et al. 2012a; Matsuda et al. 2012b; Yeager 1990b). Thus, social bonds between females may be driven by factors other than relatedness. A notable feature of social interactions in colobine monkeys, including proboscis monkeys, is non-mothers handling infants, i.e., allomothering behavior (Davies and Oates 1994; Matsuda et al. 2022; Matsuda et al. 2012b), although sometimes handlers hurt or abuse the infant and do not provide genuine care (McKenna 1979). Social bonds between females may be enhanced through such allomothering behaviors (Matsuda et al. 2015; Zhang et al. 2012), which can have various functional benefits, e.g., increased foraging efficiency for mothers and parental training of nulliparous females (Maestripieri 1994). Therefore strong ties between females within groups may develop even in the absence of close relatedness (McKenna 1979).

*F*_*ST*_-based analyses indicated that although females randomly migrated short or long distances, males were more likely to migrate relatively shorter distances than females and rarely migrated long distances, such as from the mouth of the river to >4,000 m upstream (Table 1). This may simply indicate that, in the two different bands, males disperse more proximally within the band to which they belong, whereas females can disperse proximally or distally within and out of the band.

### Inferring MLS based on matching observational and genetic data

When combining the results of the direct observations conducted in different periods from genetic research, it is more likely that females disperse more distally than males based on the results of *F*_*ST*_ analysis, which is seemingly contradictory to the results through the direct observation, i.e. the AMG utilised the riverside more extensively than OMUs, which included females (Figure 2). If a larger community structure encompassing bands A and B, which might be termed a “herd”, exists in the proboscis monkey society (Figure 4), this may explain why *F*_*st*_ was significantly different between males in the upper and lower streams. Males may disperse within a band as well as between bands, whereas females may be able to disperse beyond the herd. Because the MLS of golden snub-nosed monkeys (*Rhinopithecus roxellana*), a phylogenetically close relative of proboscis monkey, has been proposed to form grouping levels above the two-tier structure of OMUs and bands (Figure 5: Qi et al. 2014; Qi et al. 2017), the possibility of a three-tiered social organization in proboscis monkey cannot be ruled out.

**Figure 4.**
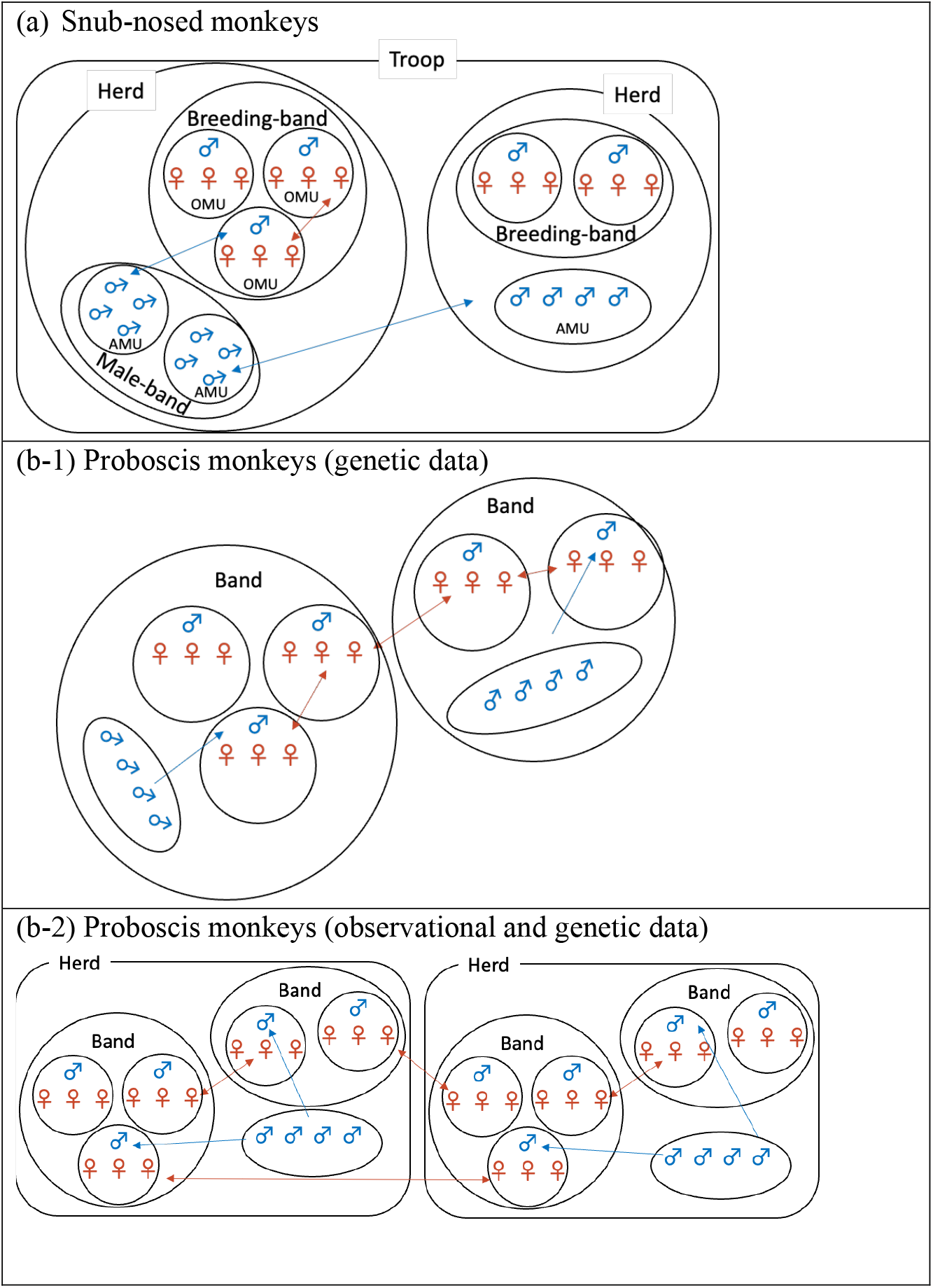
A schematic illustration of (a) a MLS in golden snub-nosed monkey based on Qi et al. (2014) where the herd level is based on spatial proximity between a band and one or more AMUs and the troop level is based on the spatial proximity between two herds; (b-1) proboscis monkey MLS based on genetic data only; and (b-2) proboscis monkey MLS inferred from observational and genetic data. Symbols and colors indicate males vs. females and lines show dispersal patterns.

The MLS of proboscis monkeys is comparable to that of snub-nosed monkeys, especially in terms of the flexible dispersal patterns seen in both sexes (Qi et al. 2009; Zhang et al. 2012). However, there is a subtle difference in that in golden snub-nosed monkeys, each breeding band composed of OMUs is accompanied by AMUs which jointly form a herd (Qi et al. 2020; Qi et al. 2014), whereas in proboscis monkeys, the AMG covers multiple bands composed of OMUs (Figure 2), although the closeness of the AMG to either band would appear to be flexibly changing over time. This inference is based on both genetic data and direct observations (Figure 4). As discussed above, results from the genetic data alone suggest that proboscis monkeys exhibit a MLS similar to that of snub-nosed monkeys. Although the lack of analysis of individual-level behavior is a limitation of this study, the more constrained dispersal patterns in males than those in females (which did not seem to migrate beyond the troop) may be a difference between the social systems of proboscis and snub-nosed monkeys. Accordingly, the social system of proboscis monkeys would be unique in that it has a patrilineal genetic basis. Assuming the MLS in proboscis monkeys is patrilineally structured, the following two observations would be well explained: 1) male–male fighting causing serious injury is rarely observed in proboscis monkeys, although sexual competition between males is intense, as evidenced by their large noses and body sizes (Koda et al. 2018; Matsuda et al. 2020b), and 2) infanticide has not been observed in natural populations of proboscis monkeys. Invoking inclusive fitness theory, if males are related to each other within a band or herd, the adult males of the OMUs are expected to tolerate each other and avoid excessive fighting over females, and likewise, the males of AMGs are expected to avoid taking over the group through serious physical challenges to the adult males of the OMUs. This aspect of the social system of proboscis monkey bears resemblance to the society of hamadryas baboons where males of the same clan (the tier above the OMU in their MLS) are philopatric, patrilineally related, spatially associated, and frequently engaged in social interactions and show mutual respect of female ‘ownership’ (Abegglen 1984; Kummer et al. 1974; Schreier and Swedell 2009; Städele et al. 2014). However, this scenario differs from that of the non-patrilineally based MLS of *R. roxellana*, where there is even collective action involving OMU males (Xiang et al. 2013).

Although infanticide has been reported in proboscis monkeys under provisioned conditions with a high population density (Agoramoorthy and Hsu 2005), infanticide has not been reported under natural conditions. Females with infants have also been reported to transfer to other OMUs, but no infanticide or attacks by dominant males have been observed in these cases (Bennett and Sebastian 1988; Matsuda et al. 2012a). After the takeover of an OMU by a male from an AMG, the risk of infanticide will be reduced if the usurping male is related to the ousted male. Hence, the mechanism underlying the lack of infanticide in proboscis monkeys may be different from that of the snub-nosed monkey, where infanticide is thought to be constrained by paternity uncertainty resulting from extra-unit mating (Qi et al. 2020). Because extra-unit mating has not been reported in proboscis monkeys (Boonratana 2011; Murai 2006; Yeager 1990a), it is unlikely that a similar scenario would act as an additional or alternative deterrent to infanticide.

Understanding why a MLS with a patrilineal basis, a key step in the evolutionary history of human society (Chapais 2008), emerged in a phylogenetically distantly related primate species, such as the proboscis monkey, may provide valuable clues about human social evolution. However, the evolutionary pathways leading to MLS in humans and colobines appear to be different; in humans, MLS are likely the product of internal fractionation of anestrally multimale-multifemale groups whereas colobine MLS seem to have emerged as a result of the amalgamation of originally autonomous OMUs (Grueter et al. 2012a). A comprehensive understanding the internal structure of multileveled societies in various animal species and inferring the process of their social evolution would further shed light on the uniqueness of human societies and contribute to developing insights into the evolutionary process of our societies.

## Acknowledgements

We express our sincere thanks to the Economic Planning Unit of the Malaysian Government, the Sabah Biodiversity Centre and the Sabah Wildlife Department for granting permission to carry out this research. We are also deeply indebted to the Sabah Forestry Department for facilitating the use of their facilities in the field. Advice and support has been generously supplied by the Higashi-laboratory members of Hokkaido University. We also appreciate the profound comments and suggestions from the members of “Tsuda CREST”. We are grateful to T. Ikeda and E. Inoue for their helpful advice on statistical analysis and genetic experiments/analyses, respectively. Finally, we are grateful to associate editor and two anonymous reviewers for their fruitful comments.

## Author contribution

I.M., T.M., B.G. and M.S. conceptualized the initial idea; I.M. and T.M. collected samples for genetic analysis and behavioural observation data, respectively; A.T. and H.B. arranged the sampling in the field; N.K.Y. and M.S. conducted a genetic experiment. I.M., T.M. and P.O. performed and interpreted the statistical analyses; C.C.G. made a significant contribution to the discussion of interspecific comparisons; I.M., C.C.G., P.O. and M.S. drafted the manuscript. All authors contributed to the final version of the manuscript.

## Declarations

### Ethics approval

the animal study was reviewed and approved by the Economic Planning Unit of the Malaysian Government and the Sabah Biodiversity Centre and was conducted in compliance with the animal care regulations and laws of Malaysian.

### Data availabilityity

All data needed to evaluate the conclusions in the paper are present in the paper and/or the supplementary materials. The data providing additional information related to this paper may be requested from the authors.

---

**Supplementary Table 1.**
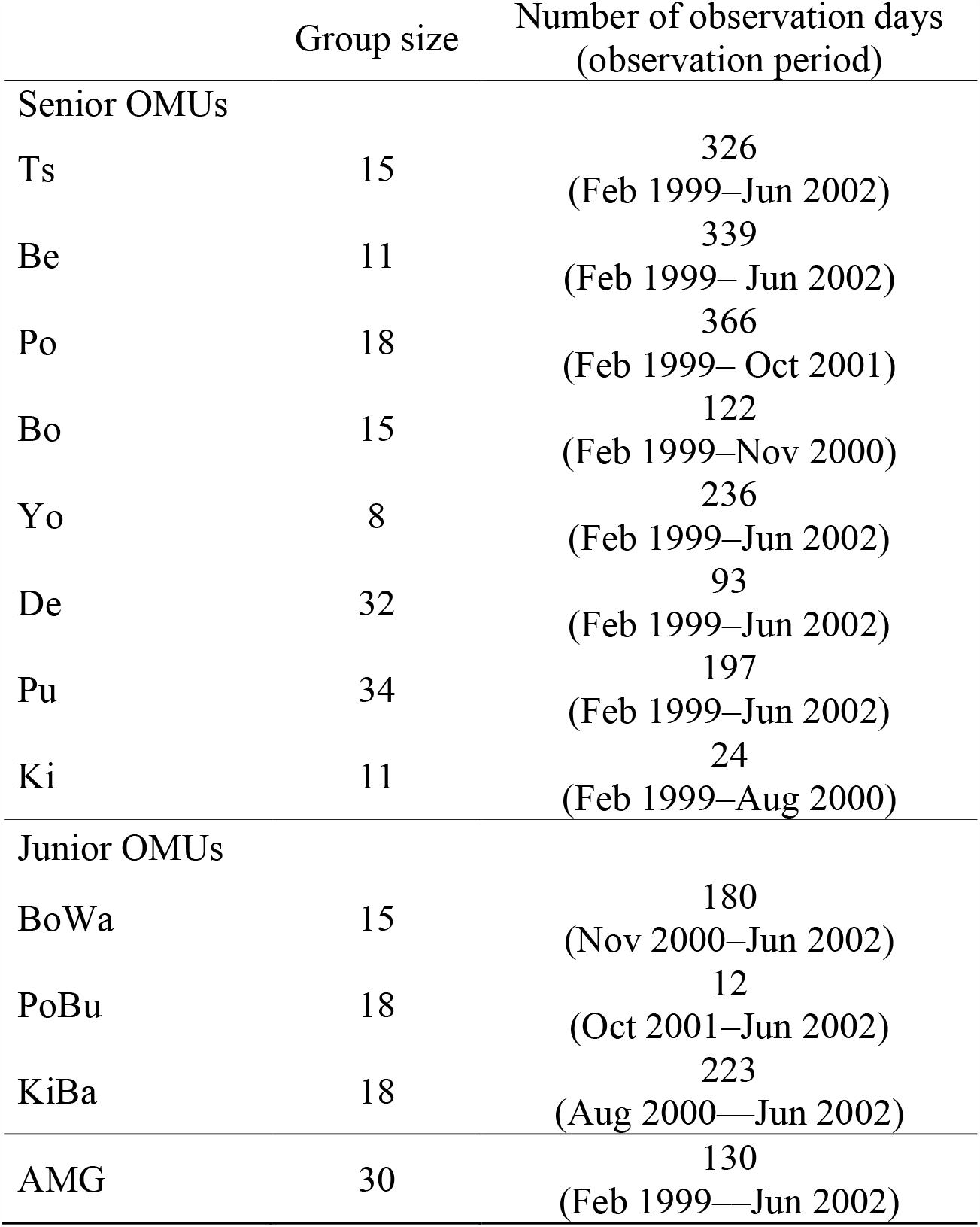
Group size and number of observation days, i.e. sleeping sites of each group were discovered.

**Supplementary Table 2.** Microsatellite marker multiplexes optimized for this study.

| **Multiplex Combination** | **Annealing Temperature (°C)** | **Markers used** | **Range** | **Dye** |
| --- | --- | --- | --- | --- |
| A | 60 | NIE10 | 165-177 | FAM |
|  |  | NIP1C5 | 197-207 | HEX |
|  |  | NIP2B8 | 229-243 | HEX |
|  |  | NIP3B6 | 238-258 | FAM |
|  |  | NIP1C8 | 220-226 | FAM |
|  |  | NIP2C12 | 187-195 | FAM |
|  |  | DEADBOX | 182-212 | TAM |
| B | 60 | NIP3C11 | 194-201 | FAM |
|  |  | NIP2D6, | 151-165 | FAM |
|  |  | NIP3B4 | 184-190 | HEX |
|  |  | NID10 | 183-185 | TAM |
|  |  | NIP3H5 | 161-169 | HEX |
|  |  | NIP4B2 | 201/203 | HEX |
|  |  | NIP2B9 | 210-220 | FAM |
| C | 62 | NIP2F3 | 175-187 | FAM |
|  |  | NIP3G2 | 232-254 | FAM |
|  |  | NIP4C11 | 245-263 | TAM |
|  |  | NIP4B1 | 152-156 | HEX |
|  |  | NIP4E10 | 207-223 | HEX |
|  |  | NIP2F7 | 138-152 | FAM |
| D | 62 | NIP1A6 | 153-159 | FAM |
|  |  | NIP2C5 | 179-187 | FAM |
|  |  | NIP3B2 | 159-171 | HEX |
|  |  | NIP3E8 | 209-233 | HEX |

**Supplementary Table 3.** Summary statistics of genetic variation between male and female proboscis monkey populations in the Menanggul River. N: sample size, Na: number of alleles, Nea: number of effective alleles, Ho: observed heterozygosity, He: expected heterozygosity, F: inbreeding coefficient, HWE: Hardy–Weinberg Equilibrium, *: significant, NS: not significant, SE: standard error for the population mean and NA: not applicable.

| Males |  |  |  |  |  |  |  |
| --- | --- | --- | --- | --- | --- | --- | --- |
| Locus | N | Na | Nea | Ho | He | F | HWE |
| NIE10 | 59 | 7.000 | 2.437 | 0.373 | 0.590 | 0.368 | * |
| NIP2C12 | 64 | 5.000 | 2.167 | 0.250 | 0.538 | 0.536 | * |
| NIP3B6 | 63 | 6.000 | 3.202 | 0.587 | 0.688 | 0.146 | * |
| NIP1C5 | 60 | 4.000 | 1.860 | 0.333 | 0.462 | 0.279 | * |
| NIP2B8 | 60 | 5.000 | 3.668 | 0.517 | 0.727 | 0.290 | * |
| DEADBOX | 64 | 2.000 | 1.969 | 0.844 | 0.492 | -0.714 | * |
| NIP2D6 | 61 | 8.000 | 3.474 | 0.492 | 0.712 | 0.309 | * |
| NIP3C11 | 62 | 4.000 | 2.133 | 0.210 | 0.531 | 0.605 | * |
| NIP2B9 | 54 | 7.000 | 4.893 | 0.463 | 0.796 | 0.418 | * |
| NIP3H5 | 60 | 4.000 | 2.131 | 0.467 | 0.531 | 0.121 | NS |
| NIP3B4 | 62 | 3.000 | 1.752 | 0.129 | 0.429 | 0.699 | * |
| NIP4B2 | 58 | 3.000 | 1.496 | 0.017 | 0.332 | 0.948 | * |
| NID10 | 60 | 1.000 | 1.000 | 0.000 | 0.000 | NA | NA |
| NIP2F7 | 63 | 3.000 | 2.881 | 0.810 | 0.653 | -0.240 | NS |
| NIP2F3 | 60 | 3.000 | 2.799 | 0.250 | 0.643 | 0.611 | * |
| NIP3G2 | 61 | 10.000 | 5.066 | 0.574 | 0.803 | 0.285 | * |
| NIP4B1 | 64 | 2.000 | 1.133 | 0.063 | 0.117 | 0.467 | * |
| NIP4E10 | 62 | 5.000 | 4.406 | 0.661 | 0.773 | 0.145 | * |
| NIP4C11 | 62 | 9.000 | 5.559 | 0.726 | 0.820 | 0.115 | * |
| NIP1A6 | 64 | 2.000 | 1.771 | 0.422 | 0.435 | 0.031 | Ns |
| NIP2C5 | 64 | 4.000 | 2.787 | 0.656 | 0.641 | -0.023 | NS |
| NIP3B2 | 64 | 5.000 | 2.260 | 0.484 | 0.557 | 0.131 | NS |
| NIP3E8 | 60 | 5.000 | 3.952 | 0.650 | 0.747 | 0.130 | NS |
| Mean | 61.348 | 4.652 | 2.817 | 0.434 | 0.566 | 0.257 | * |
| SE | 0.509 | 0.485 | 0.265 | 0.051 | 0.044 | 0.072 | NA |
| Females |  |  |  |  |  |  |  |
| Locus | N | Na | Ne | Ho | He | F | HWE |
| NIE10 | 130 | 6.000 | 3.619 | 0.262 | 0.724 | 0.639 | * |
| NIP2C12 | 129 | 6.000 | 2.413 | 0.233 | 0.586 | 0.603 | * |
| NIP3B6 | 128 | 7.000 | 4.118 | 0.477 | 0.757 | 0.371 | * |
| NIP1C5 | 122 | 6.000 | 1.957 | 0.385 | 0.489 | 0.212 | * |
| NIP2B8 | 127 | 5.000 | 2.868 | 0.299 | 0.651 | 0.541 | * |
| DEADBOX | 133 | 2.000 | 1.062 | 0.000 | 0.058 | 1.000 | * |
| NIP2D6 | 126 | 6.000 | 2.782 | 0.421 | 0.640 | 0.343 | * |
| NIP3C11 | 125 | 4.000 | 1.652 | 0.240 | 0.395 | 0.392 | * |
| NIP2B9 | 100 | 7.000 | 4.311 | 0.270 | 0.768 | 0.648 | * |
| NIP3H5 | 128 | 4.000 | 1.971 | 0.320 | 0.493 | 0.350 | * |
| **NIP3B4** | 132 | 4.000 | 1.272 | 0.114 | 0.214 | 0.468 | * |
| **NIP4B2** | 126 | 2.000 | 1.083 | 0.000 | 0.076 | 1.000 | * |
| **NID10** | 125 | 1.000 | 1.000 | 0.000 | 0.000 | NA | NA |
| **NIP2F7** | 127 | 3.000 | 2.674 | 0.693 | 0.626 | -0.107 | * |
| **NIP2F3** | 127 | 7.000 | 2.790 | 0.181 | 0.642 | 0.718 | * |
| **NIP3G2** | 119 | 10.000 | 4.313 | 0.521 | 0.768 | 0.322 | * |
| **NIP4B1** | 123 | 2.000 | 1.185 | 0.057 | 0.156 | 0.636 | * |
| **NIP4E10** | 124 | 8.000 | 4.630 | 0.508 | 0.784 | 0.352 | * |
| **NIP4C11** | 120 | 9.000 | 5.791 | 0.650 | 0.827 | 0.214 | * |
| **NIP1A6** | 133 | 2.000 | 1.975 | 0.361 | 0.494 | 0.269 | * |
| **NIP2C5** | 132 | 5.000 | 3.329 | 0.644 | 0.700 | 0.080 | * |
| **NIP3B2** | 132 | 5.000 | 2.549 | 0.447 | 0.608 | 0.265 | * |
| **NIP3E8** | 127 | 5.000 | 3.737 | 0.504 | 0.732 | 0.312 | * |
| **Mean** | 125.870 | 5.043 | 2.743 | 0.330 | 0.530 | 0.438 | * |
| **SE** | 1.430 | 0.497 | 0.275 | 0.044 | 0.054 | 0.056 | NA |

**Supplementary Figure 1.**
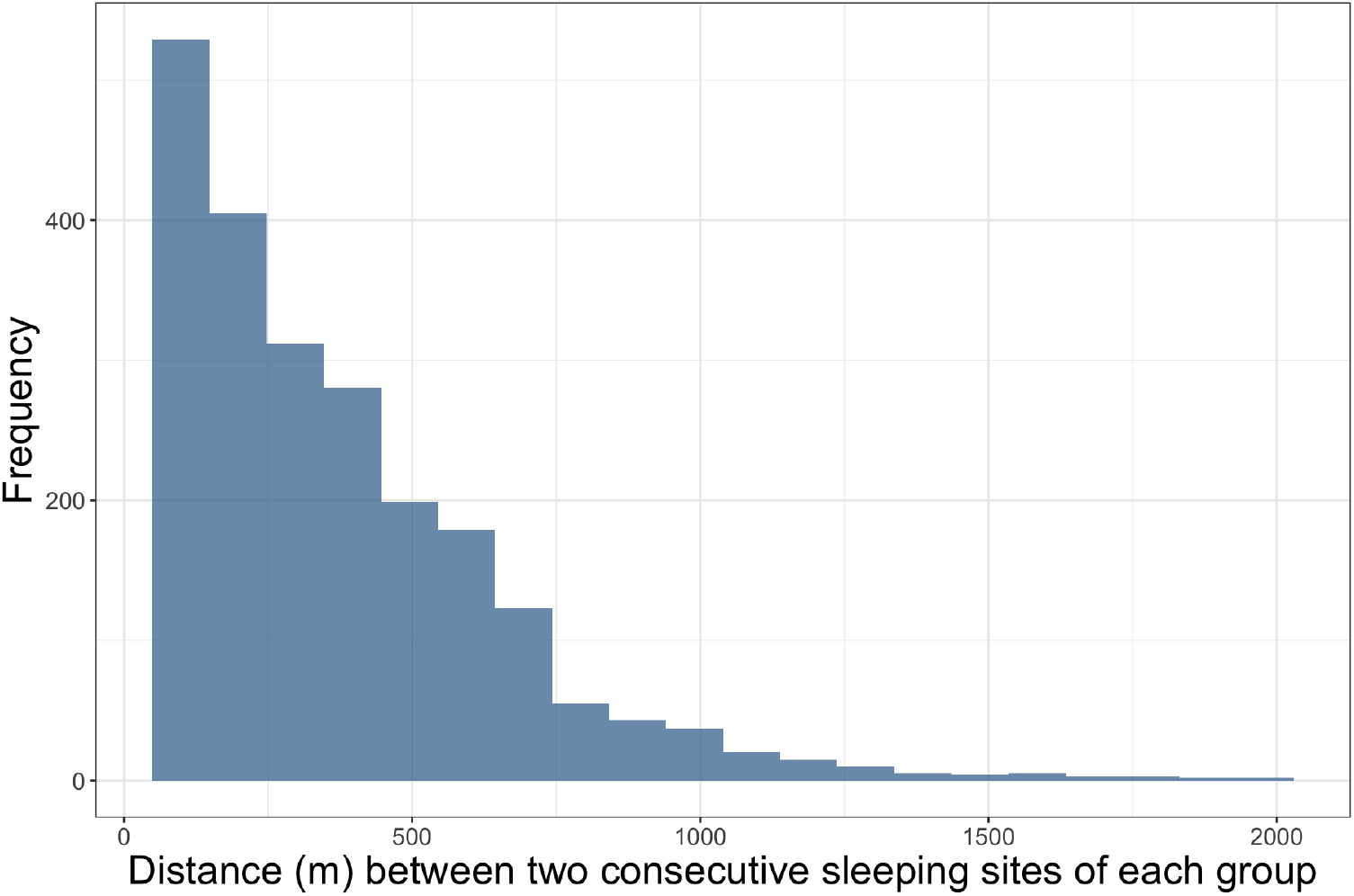
Frequency distribution of distance between two consecutive sleeping sites of each group during the study period.

**Supplementary Figure 2.**
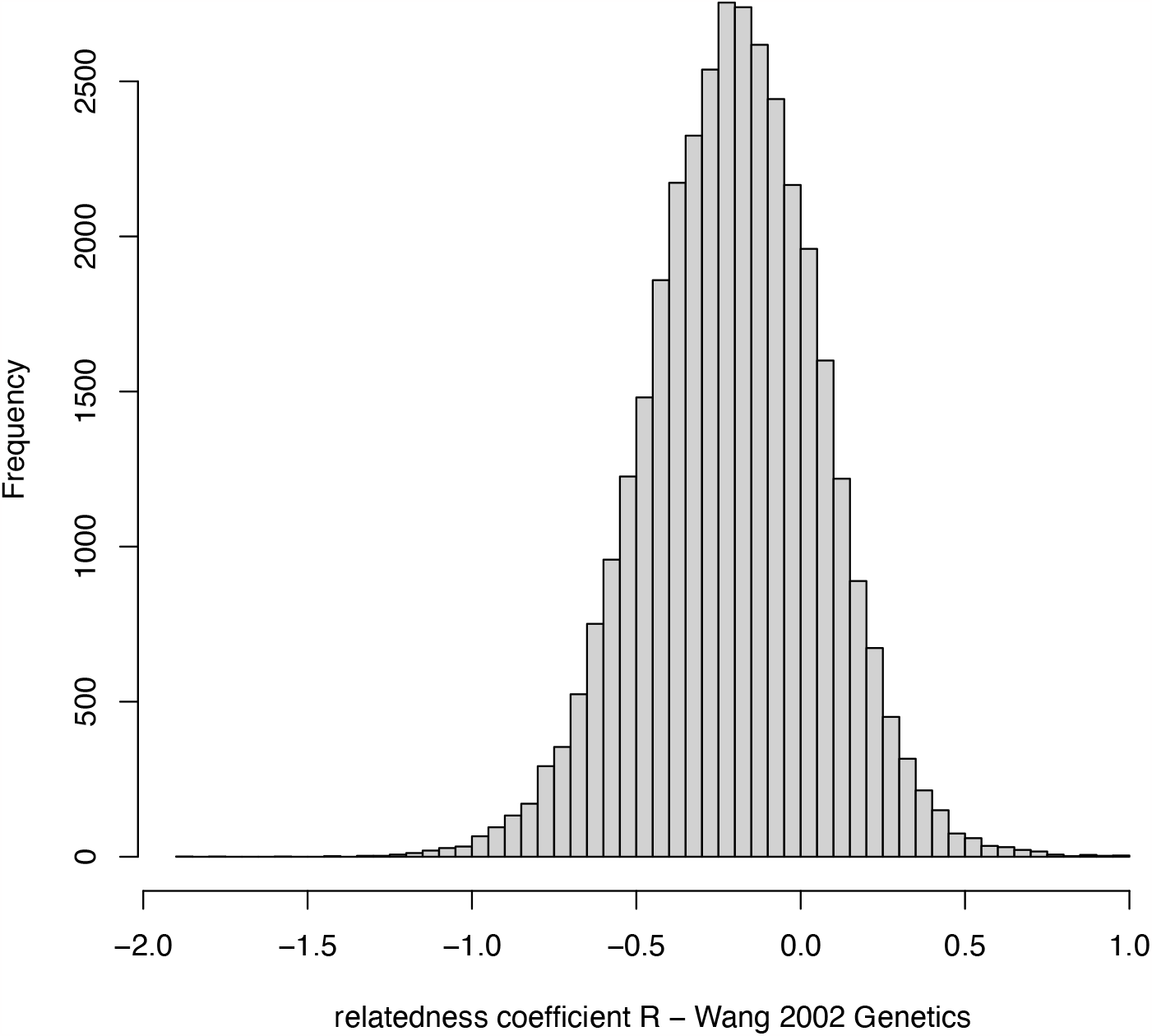
Distribution of relatedness pairwise comparisons measured among the 267 samples.

## References

Abegglen JJ (1984) On Socialization in Hamadryas Baboons: A Field Study. Bucknell University Press, Lewisburg

Agoramoorthy G, Hsu MJ (2005) Occurrence of infanticide among wild proboscis monkeys (Nasalis larvatus) in Sabah, Northern Borneo. Folia Primatol (Basel) 76:177–179. doi: 10.1159/000084380

Benjamini Y, Yekutieli D (2001) The control of the false discovery rate in multiple testing under dependency. The Annals of Statistics 29:1165–1188. doi: 10.2307/2674075

Bennett EL, Sebastian AC (1988) Social organization and ecology of proboscis monkeys (Nasalis larvatus) in mixed coastal forest in Sarawak. International Journal of Primatology 9:233–255. doi: 10.1007/bf02737402

Boonratana R (2002) Social organisation of proboscis monkeys (Nasalis larvatus) in the lower Kinabatangan, Sabah, Malaysia Malayan Nature Journal 56:57–75

Boonratana R (2011) Observations on the sexual behaviour and birth seasonality of proboscis monkey (Nasalis larvatus) along the lower Kinabatangan river, northern Borneo. Asian Primates Journal 2:2–7

Chapais B (2008) Primeval kinship: How pair-bonding gave birth to human society. Harvard University Press, Cambridge

Csardi G, Nepusz T (2006) The igraph software package for complex network research. InterJournal Complex Systems:1695

Csillery K, Johnson T, Beraldi D, Clutton-Brock T, Coltman D, Hansson B, Spong G, Pemberton JM (2006) Performance of marker-based relatedness estimators in natural populations of outbred vertebrates. Genetics 173:2091–2101. doi: 10.1534/genetics.106.057331

Davies AG, Oates FJ (1994) Colobine Monkeys: Their Ecology, Behaviour and Evolution. Cambridge University Press, Cambridge

Dunbar R, Dunbar E (1975) Social dynamics of gelada baboons. Karger, Basel

Dunbar RIM (1979) Structure of gelada baboon reproductive units I. stability of social relationships. Behaviour 69:72–87. doi: 10.1163/156853979x00403

Dyble M, Thompson J, Smith D, Salali GD, Chaudhary N, Page AE, Vinicuis L, Mace R, Migliano AB (2016) Networks of food sharing reveal the functional significance of multilevel sociality in two hunter-gatherer groups. Curr Biol 26:2017–2021. doi: 10.1016/j.cub.2016.05.064

Fang G, Jiao H-T, Wang M-Y, Huang P-Z, Liu X-M, Qi X-G, Li B-G (2022) Female demographic changes contribute to the maintenance of social stability within a primate multilevel society. Animal Behaviour 192:101–108. doi: 10.1016/j.anbehav.2022.07.018

Goffe AS, Zinner D, Fischer J (2016) Sex and friendship in a multilevel society: behavioural patterns and associations between female and male Guinea baboons. Behav Ecol Sociobiol 70:323–336. doi: 10.1007/s00265-015-2050-6

Goudet J (2005) Hierfstat, a package for r to compute and test hierarchical F-statistics. Molecular Ecology Notes 5:184–186. doi: 10.1111/j.1471-8286.2004.00828.x

Grueter CC (2022) Causes and consequences of the formation of multilevel societies in colobines. In: Matsuda I, Grueter Cyril C, Teichroeb JA (eds) The Colobines: Natural History, Behaviour and Ecological Diversity. Cambridge University Press, Cambridge, pp 293–311

Grueter CC, Chapais B, Zinner D (2012a) Evolution of multilevel social systems in nonhuman primates and humans. International Journal of Primatology 33:1002–1037. doi: 10.1007/s10764-012-9618-z

Grueter CC, Erb WM, Ulibarri LR, Matsuda I (2022) Ecology and behaviour of odd-nosed colobines. In: Matsuda I, Grueter Cyril C, Teichroeb Julie A (eds) The Colobines: Natural History, Behaviour and Ecological Diversity. Cambridge University Press, Cambridge, pp 156–185

Grueter CC, Matsuda I, Zhang P, Zinner D (2012b) Multilevel societies in primates and other mammals: Introduction to the special issue. International journal of primatology 33:993–1001. doi: 10.1007/s10764-012-9614-3

Grueter CC, Qi X, Li B, Li M (2017) Multilevel societies. Current Biology 27:R984–R986. doi: 10.1016/j.cub.2017.06.063

Grueter CC, Qi X, Zinner D, Bergman T, Li M, Xiang Z, Zhu P, Migliano AB, Miller A, Krützen M, Fischer J, Rubenstein DI, Vidya TNC, Li B, Cantor M, Swedell L (2020) Multilevel organisation of animal sociality. Trends in Ecology & Evolution. doi: 10.1016/j.tree.2020.05.003

Grueter CC, van Schaik CP (2009) Evolutionary determinants of modular societies in colobines. Behavioral Ecology 21:63–71. doi: 10.1093/beheco/arp149

Grueter CC, Wilson ML (2021) Do we need to reclassify the social systems of gregarious apes? Evol Anthropol 30:316–326. doi: 10.1002/evan.21919

Guo S, Huang K, Ji W, Garber PA, Li B (2015) The role of kinship in the formation of a primate multilevel society. Am J Phys Anthropol 156:606–613. doi: 10.1002/ajpa.22677

Hagell S, Whipple AV, Chambers CL (2013) Population genetic patterns among social groups of the endangered Central American spider monkey (Ateles geoffroyi) in a human-dominated landscape. Ecol Evol 3:1388–1399. doi: 10.1002/ece3.547

Inoue E, Ogata M, Seino S, Matsuda I (2016) Sex identification and efficient microsatellite genotyping using fecal DNA in proboscis monkeys (Nasalis larvatus). Mammal Study 41:141–148. doi: 10.3106/041.041.0304

Kawabe M, Mano T (1972) Ecology and behavior of the wild proboscis monkey, Nasalis larvatus (Wurmb), in Sabah, Malaysia. Primates 13:213–227. doi: 10.1007/bf01840882

Kern JA (1964) Observations on the habits of the proboscis monkey, Nasalis larvatus (Wurmb), made in the Brunei Bay area, Borneo. Zoologica : scientific contributions of the New York Zoological Society 49:183–192

Koda H, Murai T, Tuuga A, Goossens B, Nathan S, Stark DJ, Ramirez DAR, Sha JCM, Osman I, Sipangkui R, Seino S, Matsuda I (2018) Nasalization by Nasalis larvatus: Larger noses audiovisually advertise conspecifics in proboscis monkeys. Science Advances 4:eaaq0250. doi: 10.1126/sciadv.aaq0250

Kummer H (1968) Social Organization of Hamadryas Baboons. The University of Chicago Press, Basel

Kummer H, Gotz W, Angst W (1974) Triadic differentiation: an inhibitory process protecting pair bonds in baboons. Behaviour 49:62–87. doi: 10.1163/156853974x00408

Layton R, O’Hara S (2010) Human Social Evolution: A Comparison of Hunter-gatherer and Chimpanzee Social Organization. In: Dunbar RIM, Gamble C, Gowlett J (eds) Social Brain, Distributed Mind. Oxford University Press, Oxford, pp 83–114

Longmire JL, Maltbie M, Baker RJ (1997) Use of ‘Lysis Buffer’ in DNA isolation and its implication for museum collections. Occasional Papers, The Museum, Texas Tech University:1-3

Macdonald DW (1982) Notes on the size and composition of groups of proboscis monkey, Nasalis larvatus. Folia Primatologica 37:95–98. doi: 10.1159/000156022

Maestripieri D (1994) Social structure, infant handling, and mothering styles in group-living old world monkeys. International Journal of Primatology 15:531–553. doi: 10.1007/bf02735970

Matsuda I, Abram NK, Stark DJ, Sha JCM, Ancrenaz M, Goossens B, Lackman I, Tuuga A, Kubo T (2020a) Population dynamics of the proboscis monkey Nasalis larvatus in the Lower Kinabatangan, Sabah, Borneo, Malaysia. Oryx 54:583–590. doi: 10.1017/s0030605318000467

Matsuda I, Fukaya K, Pasquaretta C, Sueur C (2015) Factors influencing grooming social networks: insights from comparisons of colobines with different dispersal patterns. In: Furuichi T, Yamagiwa J, Aureli F (eds) Dispersing Primate Females. Springer, pp 231–254

Matsuda I, Grueter CC, Teichroeb JA (2022) The Colobines: Natural History, Behaviour and Ecological Diversity. Cambridge University Press, Cambridge

Matsuda I, Kubo T, Tuuga A, Higashi S (2010a) A Bayesian analysis of the temporal change of local density of proboscis monkeys: implications for environmental effects on a multilevel society. American Journal of Physical Anthropology 142:235–245. doi: 10.1002/ajpa.21218

Matsuda I, Nakabayashi M, Otani Y, Yap SW, Tuuga A, Wong A, Bernard H, Wich SA, Kubo T (2019) Comparison of plant diversity and phenology of riverine and mangrove forests with those of the dryland forest in Sabah, Borneo, Malaysia. In: Nowak K, Barnett AA, Matsuda I (eds) Primates in Flooded Habitats: Ecology and Conservation. Cambridge University Press, Cambridge, pp 15–28

Matsuda I, Stark DJ, Saldivar DAR, Tuuga A, Nathan SKSS, Goossens B, van Schaik CP, Koda H (2020b) Large male proboscis monkeys have larger noses but smaller canines. Communications Biology 3. doi: 10.1038/s42003-020-01245-0

Matsuda I, Tuuga A, Bernard H, Furuichi T (2012a) Inter-individual relationships in proboscis monkeys: a preliminary comparison with other non-human primates. Primates 53:13–23. doi: 10.1007/s10329-011-0259-1

Matsuda I, Tuuga A, Higashi S (2009) Ranging behavior of proboscis monkeys in a riverine forest with special reference to ranging in inland forest. International Journal of Primatology 30:313–325. doi: 10.1007/s10764-009-9344-3

Matsuda I, Tuuga A, Higashi S (2010b) Effects of water level on sleeping-site selection and inter-group association in proboscis monkeys: why do they sleep alone inland on flooded days? Ecological Research 25:475–482. doi: 10.1007/s11284-009-0677-3

Matsuda I, Zhang P, Swedell L, Mori U, Tuuga A, Bernard H, Sueur C (2012b) Comparisons of intraunit relationships in nonhuman primates living in multilevel social systems. Int J Primatol 33:1038–1053. doi: 10.1007/s10764-012-9616-1

Mbora DNM, McPeek MA (2014) How monkeys see a forest: genetic variation and population genetic structure of two forest primates. Conservation Genetics 16:559–569. doi: 10.1007/s10592-014-0680-2

McKenna JJ (1979) The evolution of allomothering behavior among colobine monkeys: Function and opportunism in evolution. American Anthropologist 81:818–840. doi: 10.1525/aa.1979.81.4.02a00040

Miller A, Uddin S, Judge DS, Kaplin BA, Ndayishimiye D, Uwingeneye G, Grueter CC (2020) Spatiotemporal association patterns in a supergroup of Rwenzori black-and-white colobus (Colobus angolensis ruwenzorii) are consistent with a multilevel society. American Journal of Primatology:e23127. doi: 10.1002/ajp.23127

Morrison RE, Groenenberg M, Breuer T, Manguette ML, Walsh PD (2019) Hierarchical social modularity in gorillas. Proc Biol Sci 286:20190681. doi: 10.1098/rspb.2019.0681

Munshi-South J, Bernard H (2011) Genetic diversity and distinctiveness of the proboscis monkeys (Nasalis larvatus) of the Klias Peninsula, Sabah, Malaysia. J Hered 102:342–346. doi: 10.1093/jhered/esr013

Murai T (2004) Social behaviors of all-male proboscis monkeys when joined by females. Ecological Research 19:451–454. doi: 10.1111/j.1440-1703.2004.00656.x

Murai T (2006) Mating behaviors of the proboscis monkey (Nasalis larvatus). Am J Primatol 68:832–837. doi: 10.1002/ajp.20266

Murai T, Mohamed M, Bernard H, Mahedi PA, Saburi R, Higashi S (2007) Female transfer between one-male groups of proboscis monkey (Nasalis larvatus). Primates 48:117–121. doi: 10.1007/s10329-006-0005-2

Ogata M, Seino S (2014) Genetic analysis of captive proboscis monkeys. Zoo Biol. doi: 10.1002/zoo.21176

Oklander LI, Mino CI, Fernandez G, Caputo M, Corach D (2017) Genetic structure in the southernmost populations of black-and-gold howler monkeys (Alouatta caraya) and its conservation implications. PLoS One 12:e0185867. doi: 10.1371/journal.pone.0185867

Patzelt A, Kopp GH, Ndao I, Kalbitzer U, Zinner D, Fischer J (2014) Male tolerance and male-male bonds in a multilevel primate society. Proc Natl Acad Sci U S A. doi: 10.1073/pnas.1405811111

Qi X-G, Grueter CC, Fang G, Huang P-Z, Zhang J, Duan Y-M, Huang Z-P, Garber PA, Li B-G (2020) Multilevel societies facilitate infanticide avoidance through increased extrapair matings. Animal Behaviour. doi: 10.1016/j.anbehav.2019.12.014

Qi XG, Garber PA, Ji W, Huang ZP, Huang K, Zhang P, Guo ST, Wang XW, He G, Zhang P, Li BG (2014) Satellite telemetry and social modeling offer new insights into the origin of primate multilevel societies. Nat Commun 5:5296. doi: 10.1038/ncomms6296

Qi XG, Huang K, Fang G, Grueter CC, Dunn DW, Li YL, Ji W, Wang XY, Wang RT, Garber PA, Li BG (2017) Male cooperation for breeding opportunities contributes to the evolution of multilevel societies. Proc Biol Sci 284. doi: 10.1098/rspb.2017.1480

Qi XG, Li BG, Garber PA, Ji W, Watanabe K (2009) Social dynamics of the golden snub-nosed monkey (Rhinopithecus roxellana): female transfer and one-male unit succession. Am J Primatol 71:670–679. doi: 10.1002/ajp.20702

Quinn GP, Keough MJ (2002) Experimental design and data analysis for biologists. Cambridge University Press, Cambridge

R-Core-Development-Team (2023) R: a language and environment for statistical computing. Foundation for Statistical Computing, Vienna

Raymond M, Rousset F (1995) GENEPOP (version 1.2): population genetics software for exact tests and ecumenicism. Journal of Heredity 86:248–249

Rodseth L, Wrangham RW, Harrigan AM, Smuts BB, Dare R, Fox R, King BJ, Lee PC, Foley RA, Muller JC, Otterbein KF, Strier KB, Turke PW, Wolpoff MH (1991) The human community as a primate society [and Comments]. Current Anthropology 32:221–254. doi: 10.1086/203952

Rousseeuw PJ (1987) Silhouettes: A graphical aid to the interpretation and validation of cluster analysis. Journal of Computational and Applied Mathematics 20:53–65. doi: 10.1016/0377-0427(87)90125-7

Rousset F (2008) Genepop’007: a complete re-implementation of the genepop software for Windows and Linux. Mol Ecol Resour 8:103–106. doi: 10.1111/j.1471-8286.2007.01931.x

Salgado-Lynn M, Stanton DWG, Sakong R, Cable J, Goossens B, Bruford MW (2010) Microsatellite markers for the proboscis monkey (Nasalis larvatus). Conservation Genetics Resources 2:159–163. doi: 10.1007/s12686-010-9295-1

Salter RE, MacKenzie NA, Nightingale N, Aken KM, Chai P. K P (1985) Habitat use, ranging behaviour, and food habits of the proboscis monkey, Nasalis larvatus (van Wurmb), in Sarawak. Primates 26:436–451. doi: 10.1007/bf02382458

Schreier AL, Swedell L (2009) The fourth level of social structure in a multi-level society: ecological and social functions of clans in hamadryas baboons. Am J Primatol 71:948–955. doi: 10.1002/ajp.20736

Schreier AL, Swedell L (2012) Ecology and sociality in a multilevel society: ecological determinants of spatial cohesion in hamadryas baboons. Am J Phys Anthropol 148:580–588. doi: 10.1002/ajpa.22076

Städele V, Pines M, Swedell L, Vigilant L (2016) The ties that bind: Maternal kin bias in a multilevel primate society despite natal dispersal by both sexes. Am J Primatol 78:731–744. doi: 10.1002/ajp.22537

Städele V, Van Doren V, Pines M, Swedell L, Vigilant L (2014) Fine-scale genetic assessment of sex-specific dispersal patterns in a multilevel primate society. Journal of Human Evolution. doi: 10.1016/j.jhevol.2014.10.019

Stead SM, Teichroeb JA (2019) A multi-level society comprised of one-male and multi-male core units in an African colobine (Colobus angolensis ruwenzorii). PLoS One 14:e0217666. doi: 10.1371/journal.pone.0217666

Swedell L (2011) African Papionins: diversity of social organization and ecological flexibility. In: Campbell CJ, Fuentes A, Mackinnon KC, Bearder SK, Stumpf RM (eds) Primates in Perspective, 2nd edn. Oxford University Press, Oxford

Swedell L, Plummer T (2012) A Papionin multilevel society as a model for Hominin social evolution. International Journal of Primatology 33:1165–1193. doi: 10.1007/s10764-012-9600-9

Tinsley Johnson E, Snyder-Mackler N, Beehner JC, Bergman TJ (2013) Kinship and Dominance Rank Influence the Strength of Social Bonds in Female Geladas (Theropithecus gelada). International Journal of Primatology 35:288–304. doi: 10.1007/s10764-013-9733-5

Van Horn RC, Altmann J, Alberts SC (2008) Can’t get there from here: inferring kinship from pairwise genetic relatedness. Animal Behaviour 75:1173–1180. doi: 10.1016/j.anbehav.2007.08.027

Wang J (2002) An estimator for pairwise relatedness using molecular markers. Genetics 160:1203–1215. doi: 10.1093%2Fgenetics%2F160.3.1203

Weir BS, Cockerham CC (1984) Estimating F-statistics for the analysis of population structure. Evolution 38:1358–1370. doi: 10.1111/j.1558-5646.1984.tb05657.x

Wright S (1951) The genetical structure of populations. Ann Eugen 15:323–354. doi: 10.1111/j.1469-1809.1949.tb02451.x

Xiang ZF, Yang BH, Yu Y, Yao H, Grueter CC, Garber PA, Li M (2013) Males collectively defend their one-male units against bachelor males in a multi-level primate society. Am J Primatol. doi: 10.1002/ajp.22254

Yeager CP (1990a) Notes on the sexual behavior of the proboscis monkey (Nasalis larvatus). American Journal of Primatology 21:223–227. doi: 10.1002/ajp.1350210306

Yeager CP (1990b) Proboscis monkey (Nasalis larvatus) social organization: Group structure. American Journal of Primatology 20:95–106. doi: 10.1002/ajp.1350200204

Yeager CP (1991) Proboscis monkey (Nasalis larvatus) social organization: Intergroup patterns of association. American Journal of Primatology 23:73–86. doi: 10.1002/ajp.1350230202

Yeager CP (1992) Proboscis monkey (Nasalis larvatus) social organization: Nature and possible functions of intergroup patterns of association. American Journal of Primatology 26:133–137. doi: 10.1002/ajp.1350260207

Yu Y, Xiang ZF, Yao H, Grueter CC, Li M (2013) Female snub-nosed monkeys exchange grooming for sex and infant handling. PLoS One 8:e74822. doi: 10.1371/journal.pone.0074822

Zhang P, Li B-g, Qi X-g, MacIntosh AJJ, Watanabe K (2012) A proximity-based social network of a group of Sichuan snub-nosed monkeys (Rhinopithecus roxellana). International Journal of Primatology 33:1081–1095. doi: 10.1007/s10764-012-9608-1

